# Distinct representations of planned reach trajectories in human premotor and posterior parietal cortex

**DOI:** 10.1101/154385

**Authors:** Artur Pilacinski, Axel Lindner

## Abstract

Goal-directed movements of the hand are often directed straight at the target, e.g. when swatting a fly; but when drawing or avoiding obstacles, hand trajectories can also become quite complex. Studies on movement planning have largely neglected the latter case and the question of whether the same neural machinery is planning straight, saccade-like vs. complex hand trajectories. Using time-resolved fMRI during delayed response tasks we examined planning activity in human superior parietal lobule (SPL) and dorsal premotor cortex (PMd). We show that the recruitment of both areas in trajectory planning differs significantly: PMd represented both straight and complex hand trajectories while SPL only those that led straight to the target. This implies that complex and computationally demanding reach planning is governed by a frontal pathway while a parietal route could warrant an alternative and faster way to put simple plans into action.

## INTRODUCTION

Goal-directed eye saccades and hand reaches share many commonalities. Both movement types are prepared based on target and effector representations in a visual (retinal) reference frame and even the neural correlates responsible for their programming do partially overlap^1^. According to a well-established view, the motor plans for saccades are thereby defined by coding a difference vector between the current position of the eye and the desired saccade endpoint^2–5^. As there are no objects in the eye socket that would interfere with the rotation of the eyeball, such simple planning scheme seems optimal for its purpose. In many cases hand movements are executed in a similar point-to-point fashion, such as when catching a ball or swatting a fly. In these latter situations the hand movement could likewise be determined by a difference vector between target and hand^6^. However, this does not always suffice: imagine you’d like to reach for your pen, but a mug of coffee sits right between the pen and your hand. In such situation your eye could still saccade straight towards the pen while any straight accompanying hand movement aimed just at the pen would cause your hand to bump into the mug with potentially severe consequences. Therefore, to allow the hand to circumvent the obstacle, an appropriate reach trajectory needs to be programmed. It seems likely, that such ability to precisely plan hand trajectories is not only required to avoid obstacles, but it perhaps does also underlie our ability to perform the seemingly endless variety of highly-complex and skillful movements of the hand, such as drawing or handwriting.

Electrophysiological research in monkeys has yielded some important clues about where and how the planning of reach trajectories could be realized by the brain. A prominent candidate for reach trajectory planning is dorsal premotor cortex (PMd), as neurons in this brain area are not merely interested in target location or the hand-target difference vector but do represent information relevant for trajectory coding. For instance, in the presence of obstacles PMd does not only code movement plans towards the target location itself but it also represents the initial direction of movement that is needed to circumvent any obstacle^15^. Moreover, Hocherman and Wise^7^ have demonstrated, that some neurons in macaque premotor cortex (as well as primary motor cortex and supplementary motor area) exhibit firing patterns that correlate with the curvature of the trajectory of an upcoming reach. Premotor coding of reach curvature may – along with the coding of initial movement direction - support the ability to circumvent obstacle. In accordance with this interpretation, ablation of premotor cortex disables monkeys’ ability to avoid obstacles and they instead attempt to reach directly towards the target ^8^. This latter experiment not only directly supports a role of PMd in trajectory planning. It also highlights that planning of straight, direct reaches is still preserved despite PMd lesions and hence such vector-like reach planning must be (also) maintained by other brain regions.

Reach-related areas within the posterior parietal cortex (PPC), namely the parietal reach region (PRR) in the medial wall of the posterior intraparietal sulcus (IPS) of macaque monkeys and its functional human homologue in neighboring parts of superior parietal lobule (SPL), are likely substrates that could subserve this function. In fact, monkey PRR and human SPL have been demonstrated to represent reaches in terms of hand-target difference vectors^6,53^, i.e. in an optimal format for coding straight reach paths. Yet, several electrophysiological studies demonstrated that these reach planning regions in PPC may also contain trajectory-related information beyond vector coding. Note, however, that unlike to the work on PMd most of these studies focused on neural activity during reach execution^9–12^ but not on planning. A notable exception is the study of Torres and colleagues^13^, who utilized a simplified obstacle avoidance task. They demonstrated that single cells in monkey PRR modulated their activity prior to the reach whenever a barrier blocked the direct reach path. It was unclear, however, whether the modulation observed in this study truly reflected the initial reach direction or, alternatively, strategical chances in initial hand posture present during the planning stage. Taken together, previous research on reach planning in monkey posterior parietal cortex has highlighted its role in the vector-like coding of reach movements. It is unclear, however, whether it also contributes to the planning of complex trajectories.

Here we tried to reveal how trajectory information is represented prior to movement execution in reach -related areas of the human brain, namely areas SPL and PMd. Specifically, we wanted to examine how trajectory representations change when a movement plan could theoretically be constructed by just defining a vector between the initial hand position and a target as compared to situations when these difference vectors are identical but the movement paths vary. Based on the aforementioned research in macaque monkeys we expected to potentially reveal trajectory plan representations in human SPL^10–12,9,13,14^ and PMd^7,14,15^ and, possibly, in primary motor cortex^7,16^ as well as supplementary motor area^7,14^. Additionally, we assumed that the trajectory representations in SPL and PMd would likely differ depending on the type of the movement required: while PMd should contribute to the preparation of complex trajectories, SPL might be exclusively engaged in planning straight and direct paths.

## RESULTS

To address our hypotheses, we conducted two human functional magnetic resonance imaging (fMRI) experiments where subjects had to plan and execute finger reaches towards visually cued targets. Two groups of twelve and seven volunteers took part in Experiments 1 and 2, respectively. All of them were right handed, had no history of neurological disease and had normal or corrected to normal vision (see “METHODS” for details). All volunteers gave their written informed consent according to the Declaration of Helsinki prior to the experiment, and the study was approved by the local ethics committee. In Experiment 1 we varied the length of complex (curved) reach trajectories while keeping the hand-target vector constant across conditions. This experiment mimicked situations that enforce the programming of detailed trajectories (like during obstacle avoidance). In Experiment 2 we varied the distance to the target, and thus the hand-target vector, while instructing subjects to perform simple, straight reaches towards it. We hypothesized that if reach trajectories are represented by neural populations reflecting the desired path, the BOLD signal amplitude should scale with certain kinematic properties of trajectories (i.e. their length or complexity) as it would capture the increasingly larger neural populations that are recruited to represent these properties as they scale up^17^.

For the purpose of our experiments, we constructed a virtual-reality reach environment, consisting of an MR-compatible resistive touchscreen panel and a rear projection display system allowing subjects to receive visual feedback about their reaching finger position in approximate spatio-temporal correspondence with the true movement (Fig. 1A). Subjects were positioned with their head tilted forward inside the head coil to allow them to naturally look in the direction matching their fingertip position although without a direct vision of their hand.

**Figure 1.**
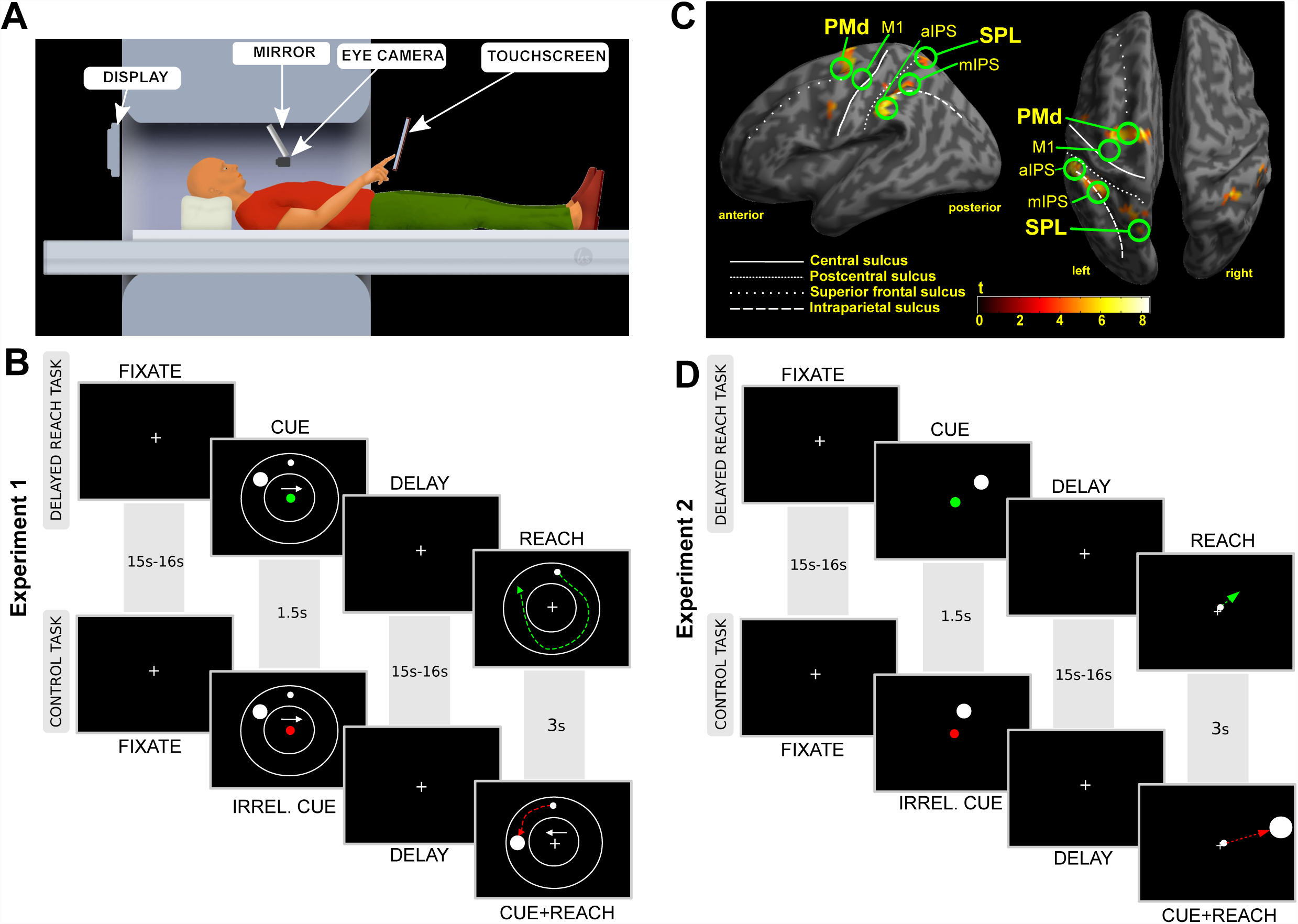
A) MRI-compatible virtual reality reach setup. Subjects viewed the rear-projected display in the mirror and didn’t have direct vision of their reaching hand. For details please refer to the main text. B) Timeline of the delayed reach task (DRT) and the control task (CT) of Experiment 1. Subjects were supposed to reach to the target (filled large white circle) by moving their finger from the starting position (filled small white circle) in either clockwise or counter-clockwise direction, as was specified by the white arrow cue. These cues were shown in the CUE period of both conditions but they were relevant only in case of the DRT. In the CT all cues were irrelevant and the ultimate movement was instructed by a new set of target and arrow cues presented during the REACH phase. Colored dashed lines illustrate putative reach trajectories in both tasks. Additional afterimage masks (500ms) presented after CUE and REACH screens are not shown (see “METHODS” for details). For better visibility the objects are plotted not to scale. C) Planning Activity. Inflated cortical surface with an overlay of the statistical contrast of delay-related planning activity (DRT>CT) obtained from 12 subjects in Experiment 1 (p<0.001, uncorrected; t-value>4.0). Labels identify regions of interest that were included in our ROI analyses. Major anatomical landmarks are labeled in addition. D) Timeline of an exemplary delayed reach (DRT) and control trial (CT) for Experiment 2 (see main text for details). See text and compare B) for detailed descriptions of the individual task phases.

### Experiment 1

The first experiment (Fig. 1B) consisted of a circular reaching paradigm comprising of two task variants: the first variant was a delayed reach task (DRT), which was used to trace reach-trajectory-related activity during planning and execution. The participants were required to remember an initially cued target location (“CUE”-phase), and then, after a delay (“DELAY”- phase), a “go” cue appeared that prompted the participants to move their finger to the now invisible target location (“REACH”-phase). The DRT was contrasted with a second task, namely a control task (CT), in which subjects’ goal was to ignore the initial spatial cue and, after a delay, to move to a visible target presented at a new location. The key difference between both tasks was that in the DRT, the subjects had to plan a movement well before its execution (during the delay epoch), whereas in the CT, the movement was only planned after the “go” cue appeared, namely when the actual target was presented. The key idea is that during the delay period of the DRT one can assess planning activity in the absence of the varying sensory cues and before a movement is being executed. By contrasting the respective activity estimates in the DRT with the CT one can further control for unspecific processes common to both tasks such as unspecific motor preparation (e.g. compare Lindner et al., 2010).

In both tasks the finger starting location was the topmost position between two large circles indicating the circular movement space. The current location of the finger was indicated by a small dot visible during the CUE and the REACH phase only. An arrow cue indicated either a clockwise (right pointing arrow) or a counter-clockwise movement (left pointing arrow) towards the target cue. Accordingly, reaches needed to be executed along a circular path of varying distance (see Fig. 1B; also compare Fig. S1 A and B). This allowed us to capture trajectory-related information and to isolate it from information related to a hand-target vector and an eye-target vector, which both were (on average) kept constant in this task. Moreover, this procedure ensured that the target and any retrospective memory thereof would be the same across conditions while reach distance (and complexity) and any related prospective processes engaged in reach planning would vary. In the CT the initial cues were irrelevant and the circular movement was specified by independently selected directional and target cues displayed during the movement epoch (Fig. 1B).

As a first step, we analyzed subjects’ behavior in Experiment 1 in terms of subjects’ reaction times as well as the duration, speed, endpoint error of movement and frequency of residual saccades. In brief, 2x2 (repeated measures) ANOVAs with the factors TASK and DISTANCE were performed on subjects’ average behavioral estimates. The respective statistical analysis of subjects’ reaction times (Fig. S1C) yielded significantly shorter reaction times in the DRT condition than in the control condition, indicating that the movements were actually pre-planned in the DRT (factor TASK: df=11, F=6.8, p=0.024, eta^2^_G_=0.0207; all other effects were not significant: DISTANCE: df=11, F=2.5, p=0.140, eta^2^_G_=0.0287; TASK*DISTANCE: df=11, F=1.5, p=0.252, eta^2^_G_=0.0041)^18^. Movement durations were significantly longer in DRT (TASK: df=11, F=27.7 p=0.0003, eta^2^_G_=0.235) and for longer trajectories (DISTANCE: df=11, F=701.4, p<0.0001, eta^2^_G_=0.908). It is noteworthy that the latter effect was driven by much larger duration differences (see Fig. S1D). There was no interaction between the two main factors (TASK*DISTANCE: df=11, F=1.7, p=0.21, eta^2^_G_=0.014). Endpoint error (see Fig. S1E) was constant across tasks and distances (TASK: df=11, F=2.4, p=0.148, eta^2^_G_=0.050; DISTANCE: df=11, F=3.5, p=0.087, eta^2^_G_=0.062; TASK*DISTANCE: df=11, F=3.1, p=0.104, eta^2^_G_=0.023). Maximal movement speed (Fig. S1F) did not differ across tasks (TASK: df=11, F=0.14, p=0.72, eta^2^_G_=0.001). It was however higher for longer trajectories (DISTANCE: df=11, F=57.87, p=0.00005, eta^2^_G_=0.589). The interaction effect was not significant (TASK*DISTANCE: df=11, F=2.71, p=0.13, eta^2^_G_=0.037). Finally, the frequency of saccades (Fig. S1G) was indistinguishable between DRT and CT in the CUE phase (TASK: df=9, F=0.14, p=0.72, eta^2^_G_=0.00061; DISTANCE: df=9, F=0.23, p=0.64, eta^2^_G_=0.00169; TASK*DISTANCE: df=9, F=1.85, p=0.21, eta^2^_G_=0.01130), and was lower for CT “FAR” reaches than in all other conditions in the DELAY phase (TASK: df=9, F=2.8, p=0.127, eta^2^_G_=0.010; DISTANCE: df=9, F=6.1, p=0.035, eta^2^_G_=0.039; TASK*DISTANCE: df=9, F=7.2, p=0.025, eta^2^_G_=0.021). Most importantly, the saccade rates in both CUE and DELAY phase of DRT did not differ for our trajectory manipulation (“NEAR” vs. “FAR”).

Subjects’ task-related brain activity was assessed by means of fMRI. Experiments were performed in a 3T Siemens Trio scanner. Functional imaging was done using EPI sequences with 2s temporal resolution and 3x3x4 mm voxel size. Functional data were analyzed using SPM8 and were modeled using a general linear model, in which we included the following regressors of interest: the main epochs of a trial (“CUE”, “DELAY”, “REACH”) were modeled separately for each experimental task (DRT vs. CT) and for each trajectory length (“NEAR” vs. “FAR”). In order to assess correlates of trajectory planning in SPL and PMd we chose a region of interest- (ROI-) based approach. In the first step we delineated a set of brain regions recruited in movement planning by contrasting delay epochs of DRT and CT. This was done by contrasting activity estimates during the delay epochs of DRT vs. CT both within the group and within in each individual. Single subjects statistical contrasts combined with anatomical criteria were used to adjust the ultimate ROI selection in order to account for inter-individual differences in functional brain organization (see “METHODS” for details).

Figures 1C and S2 depict the resulting statistical parametric map of the group analysis, exhibiting planning regions. These figures highlight our *main ROIs*, namely PMd and SPL, along with other areas engaged in motor planning, namely intraparietal sulcus (IPS), supplementary motor area (SMA). For the sake of completeness we considered these latter areas as *complementary planning ROIs.* In addition we included primary motor cortex (M1) due to its potential engagement in trajectory planning (compare ref. 7), as well as primary visual cortex (V1), which served as a *control ROI* allowing us to monitor task-unspecific brain activity reflecting visual stimulation during all trial phases. From every ROI we next extracted timecourses of BOLD-signal change throughout a trial at 1s temporal resolution. Within each individual we then separately averaged timecourses for each experimental condition. Statistical comparisons were performed across subjects’ average timecourses and between experimental conditions. Activity-timecourses were compared for trajectories of varying length/complexity and separately for each condition. Specifically, we engaged a time-resolved analysis by recruiting multiple paired t-tests performed separately for each time point. We decided for such ROI-based time-course analysis to be able to scrutinize the dynamics of activity changes in planning areas as we expected those to potentially reflect trajectory plan representations.

Figure 2A & B show respective timecourses (averaged across all subjects) that were obtained during the DRT task for both main ROIs (A: PMd; B: SPL). The leftward part of each panel depicts the timecourses aligned to CUE onset while the rightward part represents the same timecourses but aligned to the onset of the REACH-phase. Changes in planning activity in the absence of any residual CUE-related activity can be directly inspected during the late DELAY-phase (dashed boxes in Figures 2 and S3A; note that we assume a typical delay in time to peak of the even-related haemodynamic response in the human brain^19,20^, which amounts to 5-6 seconds, and a respective decay to baseline). During this time period we observed a significant increase in BOLD activity during planning of longer/more complex trajectories in PMd – as was expected (Fig. 2A; cyan shaded area). Note that this difference emerged already early after cue presentation and already then might have reflect a trajectory-related difference in planning at early stages. However, as was pointed out before, additional CUE-related modulations of the fMRI-signal can – even if unlikely – not be completely ruled out. Finally, the difference between conditions was also present during the REACH-phase. Note, however, during this period signal modulations are contaminated by systematic differences between conditions such as movement duration or speed and the related differences in visual movement feedback (as is further illustrated below). In contrast to PMd, such trajectory-related signal modulation was virtually absent in SPL (Fig. 2B). This result speaks against a major contribution of SPL to the planning of complex trajectories. Finally, the absence of any trajectory-related variation of BOLD-signals in the DELAY phase of the control task in our main ROIs suggests that the observed signal differences in PMd are not merely due to unspecific motor preparation (Fig. S3B). In the REACH-phase, however, PMd also exhibited a significantly higher signal amplitude during FAR as opposed to NEAR reaches (Fig. S3B). As was mentioned before for the DRT, this activity pattern is likely accounted for by the systematic differences in movement execution and movement feedback.

**Figure 2.**
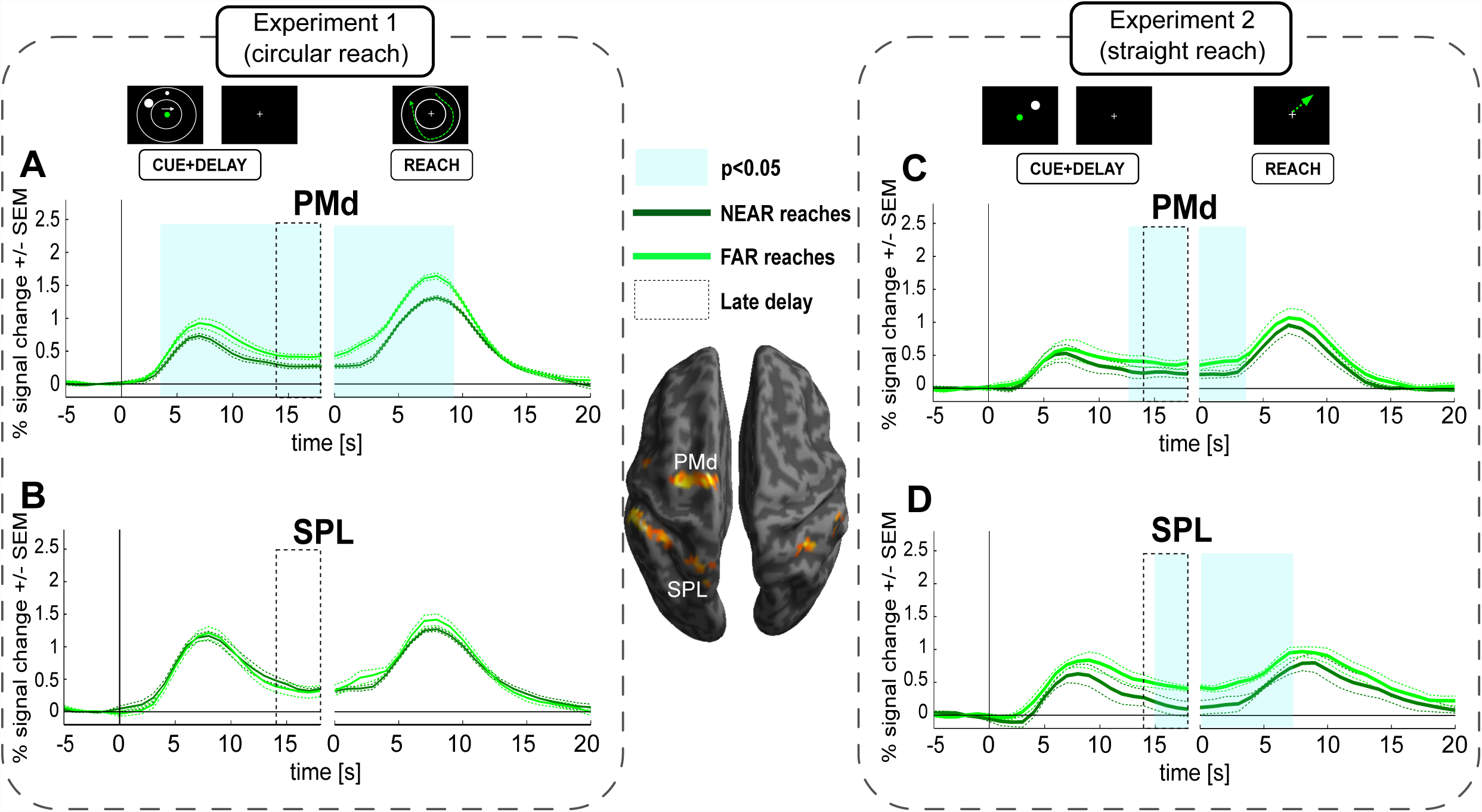
ROI timecourses extracted from PMd (A & C) and SPL (B & D), comparing the fMRI-signal in the delayed reach task of Experiments 1 and 2 (A & B vs. C & D, respectively). Cyan-shaded areas represent time epochs during which paired t-test comparisons of signal amplitudes between “NEAR” and “FAR” reaches revealed statistically significant differences (p<0.05) for at least three neighboring time-points. Such differences were considered indicative of an influence of trajectory. PMd shows different planning-related signal amplitudes in both experiments (leftward part of the panels A and C, aligned to CUE onset). SPL shows trajectory planning signal modulation in Experiment 2 only (D). PMd shows significant modulation in the reach phase in both experiments (A and C, right panels, aligned to REACH onset), whereas SPL shows such statistically significant difference only in Experiment 2 (D) although a hint of the same effect might be present in Experiment 1 as well (B). Dotted gray boxes indicate late delay phase, in which activity merely represents planning but no longer CUE-related activity. This period was also used for a subsequent statistical comparison between PMd and SPL.

In none of our additional ROIs we could reveal a significant signal-difference between NEAR and FAR during the late DELAY-phase. It is noteworthy that in M1 we also observed a significant effect of trajectory but early during the DELAY-phase (Fig. S3A). Finally, like for PMd we observed an effect of reach trajectory during reach execution in V1, M1, SMA, mIPS and aIPS (rightward panels in Fig. S3A). In all cases activity was higher for the more complex/longer trajectory. Note, however, the presence of these effects is not necessarily related to planning. It might be rather explained by the systematic differences in movement or - as is clearly indicated by V1 - by the respective amount of visual motion. As was true for PMd and SPL, we did not see any trajectory-related variation of BOLD-signals in the DELAY-phase of the control task in either of the additional ROIs, but only during the REACH-phase (see Fig. S3B).

In summary, in Experiment 1 we demonstrate that PMd – and potentially also M1 – represent plans for reach trajectories. This was evident from the fact that planning activity reflected differences in the length of curved trajectories despite the initial hand-target difference vectors were identical across trials. In the next experiment we tried to reveal potential trajectory representations during a situation in which movements were supposed to be directed straight towards a target and could thus – at least potentially – be defined by such a hand-target difference vector.

### Experiment 2

In the second experiment the overall design was similar to the one used in Experiment 1 in that we contrasted a delayed reach planning task with a direct reach task. This time, however, we used a simple center-out reaching task (compare Fig. 1D). Such task should allow us to see whether the potential trajectory-related scaling of the BOLD-signal would be seen in brain activity even if a given reach trajectory could be defined by a simple difference vector between target and hand, as such vector-based programming has been suggested based on behavioral findings^21,22^. We manipulated reach amplitude by positioning the targets at two different distances and at randomly chosen radial positions in the upper-right quadrant of the visual field (see Fig. S4A & B for examples). The idea behind this manipulation was to additionally uncover potential trajectory representations for simple, straight reach plans, while the planning of longer trajectories should result in higher BOLD signal amplitudes (compare INTRODUCTION).

Similar to Experiment 1, reaction times (Fig. S4C) were significantly shorter in the DRT (“TASK”: df=6, F=7.8, p=0.031, eta^2^_G_=0.0039) suggesting that subjects preplanned their movements in this condition. The other effects were not significant (“DISTANCE”: df=6, F=1.2, p=0.322, eta^2^_G_=0.1786; “DISTANCE*TASK”: df=6, F=1.9, p=0.220, eta^2^_G_=0.0073). As in Experiment 1, movement durations (Fig. S4D) were significantly longer for longer trajectories (“DISTANCE”: df=6, F=42.8374, p=0.00061, eta^2^_G_=0.45535). All other effects were not significant (“TASK”: df=6, F=0.5772, p=0.47619, eta^2^_G_=0.00360; “DISTANCE*TASK”: df=6, F=0.0014, p=0.97177, eta^2^_G_=0.00001). The endpoint error sizes (Fig. S4E) were significantly higher in the DRT (“TASK”:df=6, F=34.17, p=0.0011, eta^2^_G_=0.6545). This difference likely resulted from lower precision of memory- vs. visually-guided reaches. Most important for our study, both the factor distance and its interaction with task were not significant (“DISTANCE”: df=6, F=0.20, p=0.6735, eta^2^_G_=0.0045; “DISTANCE*TASK”: df=6, F=0.13, p=0.7326, eta^2^_G_=0.0045). Maximal speeds (Fig. S4F) were significantly higher for longer trajectories (“DISTANCE”: df=6, F=77.86, p=0.00012, eta^2^_G_=0.26356). All other effects were not significant (“TASK”: df=6, F=0.20, p=0.67151, eta^2^_G_=0.00043; “DISTANCE*TASK”: df=6, F=0.47, p=0.51874, eta^2^_G_=0.0014). Finally, saccade frequencies (Fig. S4G) were not different across conditions both in the CUE (“TASK”: df=4, F=1.1, p=0.358, eta^2^_G_=0.043; “DISTANCE”: df=4, F=4.8, p=0.093, eta^2^_G_=0.090; “DISTANCE*TASK”: df=4, F=4.1, p=0.114, eta^2^_G_=0.112) and in the DELAY phase (“TASK”: df=4, F=0.91, p=0.39, eta^2^_G_=0.0041; “DISTANCE”: df=4, F=0.51, p=0.51, eta^2^_G_=0.0034; “DISTANCE*TASK”: df=4, F=1.50, p=0.29, eta^2^_G_=0.0048).

We will next consider task-related changes of brain activity in our main and in the complimentary planning-related ROIs. Note that we used a similar procedure for ROI selection to the one used in Experiment 1. The actual brain regions selected for further ROI analyses were – besides, SPL and PMd - practically the same as in Experiment 1 (see figure S2, compare also Materials and Methods for further details on ROI selection).

The BOLD signals in these ROIs during the reach phase of the DRT were quite similar to those observed in Experiment 1: longer trajectories yielded larger signal amplitudes in PMd, SPL aIPS, mIPS, SMA and M1 (see Fig. 2C&D and S5A, rightward part of panels). More importantly, planning-related BOLD signals extracted during the late DELAY phase of the DRT were markedly higher for longer trajectories not only in PMd but this time in the SPL too (Fig. 2C & D; compare time period indicated by the dashed box in the leftward part of each panel). Higher delay-related BOLD signals for longer trajectories were also observed in two of our complimentary ROIs: SMA and aIPS (Fig. S5A). No planning-related signal modulation was observed in M1 or in any other additional ROI. In the control task, no ROI showed any trajectory-related activity during the DELAY phase (Fig. S5B). Only during the REACH phase, M1 and SMA exhibited a modulation of the BOLD-signal as a function of trajectory (Fig. S5B). This resembled their respective signal changes during the REACH phase in DRT (Fig. S5A) and likely can be attributed to the systematic differences in movement execution (also compare Experiment 1).

Finally, to test for the principled difference in the activation pattern in our main ROIs, SPL and PMd, across the two experiments, we performed an additional mixed model ANOVA with the factors “Experiment”, “DISTANCE” and “ROI”, comparing the activity estimates of the late delay phase of the DRT trials. These estimates captured the average activity during the last four seconds of the DELAY phase (see dashed boxes in Fig. 2 and S5). The analysis revealed a significant three-way interaction (F=6.026, df=17, p=0.025, eta^2^_G_=0.0261), further confirming that SPL and PMd exhibited diametrically distinct patterns of planning activity in both tasks, namely a (stronger) contribution of PMd to the planning of complex trajectories in the DRT of Experiment 1, while both areas represented the straight, vector-like movement trajectories in the DRT of Experiment 2.

## DISCUSSION

In Experiment 1 we showed that different reach trajectories for targets kept at the same visual locations produce differential planning responses in dorsal premotor cortex but not in SPL. Experiment 2 allowed us to further demonstrate that trajectories are represented in PMd even if reaches could, at least in principle, be coded by a simple hand-target difference vector. Moreover, we show that the activity modulated by the trajectory of straight reaches is also visible in the medial portion of SPL. Comparing the results from these two experiments, we may note that while PMd contains representations of trajectories irrespective of their complexity, SPL (and perhaps also supplementary motor area) primarily encode trajectory plans for simple reaches directed straight towards a target.

Note that we ensured the reported differences could not be accounted for by subjects’ residual eye movements (Fig. S1G & S4G). Moreover, constant error rates across conditions, as were present in both experiments (Fig. S1E & S4E), suggest that the different planning-related signals did not simply result from increasing task difficulty, but rather reflected parameters of planned trajectories. The particular design of Experiment 2 further ensured that such differences in task difficulty between “NEAR” and “FAR” should not arise in the first place (compare METHODS section). Finally, in Experiment 1 we instructed the same target locations (across conditions) while varying the way to the target (i.e. the trajectory). This allowed us not only to keep eccentricity/direction of target location balanced across conditions but this also guaranteed that any attention towards the target locations (or cues), or any retrospective memory thereof, would likewise be identical across tasks. Hence, the reported differences in brain activation should exclusively relate to the process of planning different reach trajectories. But how could such varying trajectories be realized by the brain? One possibility, suggested by prior literature, is that trajectory is initially defined by a vector pointing either towards the final target location or, alternatively, towards the initial direction of movement^15^. Then, during reach execution, the hand would be guided on-line by a feedback-based control system^22–24^. This way, trajectory would require only the first desired state (goal) to be planned in advance. As an alternative to the above, it may be hypothesized that the reach trajectory is constructed and represented as a whole at the initial stages of reach planning^17^ and only then, this initial plan is being converted to respective motor commands during movement execution while likewise allowing for on-line corrections for potential movement inaccuracies. Unfortunately, most previous research on trajectory coding concentrated on the stage of movement execution, albeit with some exceptions which (also) focused on reach planning prior to execution^13,7,15^. On the basis of the electrophysiological results provided by these latter studies, however, one could not determine whether the changes in planning activity reflect the whole trajectory of an upcoming reach or only specific factors like a change in the initial direction of the reach as enforced by additional cues (e.g. obstacles)^7,15^, or changes of the initial hand posture^13^. In the latter cases, the remaining parts of movement planning and execution could still be guided by the aforementioned on-line control system. In fact, findings of Pearce and Moran even suggested the latter possibility, as in their study the population activity in PMd seemingly encodes the initial direction of an upcoming movement regardless of the target position^15^. Our results, in turn, seem to support the idea that PMd codes the whole trajectory prior to a movement. This is because we revealed changes in planning activity for changes in overall trajectory, and despite the fact that – on average - target direction and initial movement direction were constant across varying trajectory conditions (Experiment 1). PMd activity thereby represented trajectory information even when the situation didn’t require the same precise programming of trajectories as when e.g. circumventing obstacles (Experiment 2), highlighting a vital and general role of premotor cortex in trajectory planning. Given the limitations of our experimental design and of our recording methods, however, we cannot further detail the precise nature of the trajectory parameters underlying the signal changes that we revealed. The finding of Messier and Kalaska, who reported that individual PMd neurons code both reach amplitude and direction, may present important hints^25^.

Besides PMd also posterior parietal areas exhibited activity which increased during the planning of straight, direct reaches towards more eccentric targets in Experiment 2. One might ask whether SPL thereby encoded just the difference vector or the whole straight trajectory as is defined by such vector? As suggested by some anatomical studies, SPL (and other parietal subregions) may contain a relative over-representation of the visual periphery as compared to lower visual areas^26–28^. The mere coding of the difference vector could, accordingly, recruit larger neural populations representing more peripheral targets, and potentially lead to an increase in the total BOLD signal for more eccentric target locations in Experiment 2. However, as was demonstrated by Kimmig et al. In their study on saccades^30^,the coding of larger difference vectors (or more eccentric target locations) themselves is not sufficient to modulate the amplitude of the BOLD signal in the way we observed here. Moreover, electrophysiological studies demonstrate that PPC still accentuates the central visual field over the periphery^29^, suggesting the exact opposite pattern of findings could be expected (stronger signals for more central targets) than what we described here. For these various reasons we propose an alternative explanation. In our view, the observed increase in SPL activity for more eccentric reaches in Experiment 2 is consistent with the idea that its role is to localize the target and to initially represent the trajectory in a simplified way, possibly defined along a linearly interpolated movement path that is defined by the hand-target difference vector^17^. It can then be speculated that such direct and straight trajectory representation is quasi-automatically represented by neural populations in SPL no matter what sort of movement path is actually required (such as to avoid an obstacle). This is consistent with the lack of a trajectory-related modulation of the BOLD signal amplitudes in Experiment 1, where we kept the difference vectors constant. The fact that we localized representation of such straight movements in SPL could explain why monkeys with lesions of premotor cortex cannot plan trajectories that would allow to avoid obstacles but still try to directly reach towards targets without success^8^. Moreover, our findings seem to be in accordance with the finding that PPC inactivation in monkeys caused an impairment of executing reaches along straight paths^31^. Such simplified reach paths may be useful in various everyday situations and crucial whenever a rapid response is required (e.g. when swatting a fly). Note, however, that there may still exist certain differences between the movements planned in this experiment and the fast, saccade-like hand movements performed in more natural settings.

Given the differences in movement representation between PMd and SPL as were described above, one could conceive a hierarchical model in which an initial trajectory plan is formed in SPL based on the difference vector pointing directly towards the target. As information transfer from posterior parietal cortex to M1 is faster than to PMd^32^, the simple movement plan may be quickly put into action. If required, this initial plan is “overwritten” by other frontal areas (such as PMd), which possibly consider additional information like obstacle position^13,15,33^ that would interfere with execution of the reach along the initially defined, direct path. PMd might incorporate such additional information to construct a global (potentially more complex) trajectory plan, which is passed on to areas responsible for its further processing and execution (like M1). In fact, as already described above, lesions to premotor cortex of macaque monkeys result in straight reaches, oriented directly towards targets making them unable to avoid obstacles or, alternatively, to update their initial motor plan^8^. In addition, PMd has been shown to highlight those motor plans that are actually selected for execution, rather than merely representing all the possible plans in a given context. This does further imply a role of PMd in forming the ultimate trajectory plan ^34,35^. It is worth to note, that several authors postulated that PMd may play a governing role in the sensorimotor system, modifying motor plans as needed by current context^36,37^. Such detailed hierarchy amongst the cortical areas engaged in planning reach trajectories cannot be reliably assessed on the basis of our experiments. The nature of the BOLD signal does not allow us to distinguish incoming neural signals from local processing^3^ - a distinction which is critically needed to establish hierarchy. Moreover, such distinction is perhaps particularly challenging when considering posterior parietal and premotor areas that are linked by single-synapse pathways^39–43^. To further detail how exactly trajectory information is represented and transferred throughout the network of areas engaged in sensorimotor processing, causal methods could be utilized in the future to disrupt information flow between specific regions.

In accordance with earlier studies, we did observe trajectory information encoded in M1 activity during movement execution^7,16,44^. The results revealed in Experiment 1 further suggest that M1 might encode trajectory information already at the very early stages of planning reaches along complex paths. In Experiment 2 we did not observe any M1 modulation of this kind, while in this experiment trajectories did not differ with respect to their complexity. The overall findings suggest that only trajectories of greater complexity may require engagement of M1 well before movement execution.

Another interesting finding is the involvement of supplementary motor area in the planning of straight but not of circular movements. This result parallels earlier findings of Hocherman and Wise^7^ who also reported SMA to be more active in coding straight than curved reach paths, as evident from the number of neurons responding to either of those. Hence, similar to SPL, SMA seems to be more involved in coding direct, straight trajectories. Yet, it still remains to be determined what exact role the SMA plays in this process and whether our observation can be confirmed.

In conclusion, our study shows that trajectory information is represented in premotor and posterior parietal areas of the human brain well before movement execution. Moreover, we reveal differences in the representation of planned reach trajectories across these areas. Specifically, premotor cortex can seemingly encode complex reach trajectories while posterior parietal cortex and supplementary motor area rather represent plans for movement along a simplified, straight and direct path. This parallel and distinct representation of two fundamentally different types of trajectory plans clearly asks for a meaningful functional interpretation. It is conceivable that an evolution of two disparate reach planning subsystems was desirable from an ecological point of view by offering a high degree of flexibility in adjusting hand movement control to situational demands. This way the parietal system could allow to rapidly acquire targets in a straight and simple fashion whereas the frontal system would take over whenever movements have to be performed with finesse and when the right movement path is an integral part of the motor goal.

## METHODS

### Participants

Twelve healthy, right-handed participants (11 females) in the age range of 20-32 years (mean age 25 years), participated in Experiment 1. Seven healthy, right-handed volunteers (6 females, age range 20-31 years, mean age 25 years) took part in Experiment 2. Out of these, five subjects had also participated in Experiment 1 (two of them had completed Experiment 2 first). The over-representation of female subjects in both experiments resulted from spatial constraints given our setup (especially touchscreen size and its position, see Fig. 1A), which required particularly slim subjects.

The number of participants was guided by a power analysis (power=0.80; alpha=0.05) that was informed by the descriptive statistics of a timecourse analysis on a previously published, similar fMRI dataset. In that study, planning activity varied as a function of movement sequence length^33^. For the power analysis we considered the within-subject activity difference during the late delay period (last 4 sec) in left PMd, namely for a delayed response task that required the planning of a less complex (2 targets) vs. a more complex (4 targets) movement sequence. This analysis suggested a sample size of 11 subjects (two-tailed tests). For Experiment 2 we relaxed this criterion (one-tailed tests), as we had a directional hypothesis (the stronger the activity the more complex trajectory planning). Note that here we measured each experimental condition 25 times per individual, while the study that informed our power analyses only comprised of 9 repetitions per condition.

### MR-compatible reach setup

We realized our experiments in a custom made MRI-compatible virtual reality reach setup, in which we could record 2D movements of subjects’ right index finger and could provide subjects a virtual visual representation of their finger on a stimulus screen (see Fig. 1A). Specifically, visual stimuli were projected via an LCD projector onto a translucent screen, mounted directly behind the head coil of the scanner (1024x768 pixels; 60Hz refresh rate). Subjects viewed the stimulus screen via a mirror, positioned in front of the participant. Viewing distance was approximately 82cm and roughly matched the distance from participants’ eyes to the touchscreen. To track subjects’ finger movements we used a MRI-compatible motion capture system, utilizing a resistive touchscreen panel from MAG (<u>www.magconcept.com</u>), mounted on a plastic board. This touchscreen-board was placed on top of a plastic rack onto which the stimulus-mirror, a camera for eye movement recordings and the display screen were mounted in addition. Limited by the spatial constraints of the scanner environment, we always tried to approximate a parallel alignment between the touchscreen and the display to guarantee approximate spatio-temporal correspondence between measured finger position and visual feedback thereof. Subjects were positioned with their head tilted forward inside the scanner head coil, so that they could directly look towards their pointing finger. Ultimately, direct vision of the hand was blocked by both the mirror and additional masks and subjects had to rely on the virtual visual feedback about their finger position instead. All reaches were performed in darkness and the only visual information provided was the one projected through the display system. In order to minimize potential disturbances of the magnetic field by hand and arm movements we stabilized each subject’s arm, elbow and shoulder with foam cushions and adhesive tape, so that only wrist and finger movements were made possible. To minimize movement friction we had subjects wear a cotton glove on their reaching hand.

Each of our experiments was preceded by a training session during which the subjects familiarized themselves with the tasks demands. All subjects were additionally required to practice the experiment for a minimum of 10 minutes inside the scanner once the MRI setup had been completed.

### Eye recordings

Eye fixation was monitored at 50Hz sampling rate with an MR-compatible combined camera and infra-red illumination system (MRC Systems) using the ViewPoint software (Arrington Research). Due to technical difficulties of recording eye movements in the experimental environment (extensive video capture noise, too long setup time) we were only able to perform systematic eye-recording analyses in 10 out of 12 subjects in Experiment 1 and in 5 out of 7 subjects in Experiment 2. All eye movement analyses were performed off-line using custom routines written in Matlab (MathWorks). In brief, eye position samples were filtered using a second-order 10Hz digital low-pass filter. Saccades were detected using an absolute velocity threshold (20 degrees per second), and blinks were defined as gaps in the eye position records caused by eyelid closure. Time periods with blinks were excluded from subsequent analysis. We instructed the subjects to continuously fixate on the central spot. While our subjects fulfilled this requirement in the majority of trials, we still assessed the frequency of residual saccades (amplitudes ≥ 1 deg visual angle) on a trial by trial basis and compared saccade frequencies in the CUE and in the DELAY epoch across conditions to control for potential eye movement-related confounds.

### Experiment 1

The detailed paradigms of Experiment 1 are depicted in Figure 1B. Each trial started with a 15s/16s fixation period (FIXATE), during which subjects were instructed to fixate a centrally positioned fixation cross. In addition, subjects were required to perform “finger fixation” by placing their right index finger on a tactile cue on the touchscreen. This tactile cue also defined the starting position for reaching and it would corresponded to a location at the topmost position between the circles marking the reaching space. Eye blinks were allowed though discouraged during this period. Next, a CUE screen appeared for 1.5 seconds, indicating the experimental condition (a red central cue indicating CT and a green cue indicating DRT), a target location, reach direction (an arrow indicating clockwise or counterclockwise direction), eye and finger fixation points and instructed reach space boundaries (compare Fig. 1B). Both the starting location and all targets were positioned at a constant radius of about 3deg visual angle from the fixation point. We used a predefined set of four target locations, placed in the upper-portion of the reach space either at 10 o’clock (-60°), 10.30 o’clock (-40°), 1.30 o’clock (+40°) or 2 o’clock (+60°). Note that the starting position corresponds to 0° (12 o’clock). This way, by manipulating reach direction and target location, we could alter the movement trajectory, without affecting target eccentricity and, accordingly, the hand-target difference vector. In the DRT condition subjects were required to remember the target location and to plan a movement to it according to the arrow cue. Subjects were told to ignore target and arrow cues in the CT condition, as the relevant cues would be delivered only later in the REACH phase. In both conditions subjects were asked to maintain fixation and avoid blinking during this CUE period. Next, we presented an image for 500ms, which was made up of (400) randomly positioned, black and white circles approximately the size of the cursor, to mask any after-images of the cues (not shown in Fig. 1B). This mask was followed by a DELAY period lasting 15s-16s. During the DELAY subjects were instructed to keep fixation and, again, blinking was allowed though discouraged during that period. Note that we assume that correlates of goal-directed movement planning should be present during this phase in DRT but not in CT. Finally, the response screen appeared for 3s signaling the REACH phase. In the DRT subjects had to move their right index finger to the pre-cued target location as fast and accurate as possible through a single, smooth movement of their finger. In the CT subjects were presented a new target location and a new arrow cue, and had to immediately perform a movement according to these cues. Once reaching the instructed goal, subjects had to stay at the final location until the response screen disappeared. Then, a blank screen appeared for 4s (not shown in Figure 1B) and subjects had to return their finger to the tactile cue. They were also encouraged to blink specifically during this period to reduce corneal drying in the face of prolonged periods of fixation. Note that visual feedback about finger position was only provided during the CUE and REACH phases of a trial. All experimental conditions were presented randomly interleaved and were repeated 25 times across 5 consecutive scanning sessions per subject.

### Experiment 2

The overall design of Experiment 2 was similar to Experiment 1 (compare Fig. 1D). Each trial started with a 15-16 seconds fixation epoch. The points of fixation of eye and finger overlapped spatially and corresponded to the center of the display. Then, a CUE screen was displayed for 1.5s, with a task cue presented centrally at the fixation point (a red cue indicating CT and a green cue indicating DRT), and a target cue in periphery at about 3.2deg or 7.2 deg visual angle for NEAR and FAR conditions, respectively. Target size in NEAR conditions was 0.8 deg visual angle. To accommodate for an increase in movement difficulty (ID) with increasing distance (D), we increased the size of the target (W) in the FAR conditions according to Shannon’s formulation of Fitts’ Law^50^, expressed as:

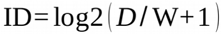

In DRT trials, subjects were instructed to remember the target cue and plan a movement to it, whereas in CT trials they were told to ignore the initial cue. The CUE screen was then masked for 500ms and a DELAY period followed, lasting 15-16 seconds. Ultimately, the REACH screen appeared for 3s and subjects had to move the cursor to the remembered target location in DRT, or to the newly cued target location in CT. After the instructed target location was reached they had to maintain their finger position at this location until the end of this task period. Then the screen was blanked and subjects had to return to the starting position. Subjects were required to perform straight movements, without lifting the finger off the touchscreen and they were told to be “as fast and as accurate as possible”. Else they did not receive any additional instructions on how to plan/perform their reaches, as we did not want to bias their natural planning strategies. As in Experiment 1, we presented all experimental conditions randomly interleaved and repeated them 25 times across 5 consecutive scanning sessions per subject.

### Finger movement analysis

Finger movement data were preprocessed using custom routines programmed in Matlab (MathWorks) and analyzed statistically using R (R Foundation for Statistical Computing). In brief, during preprocessing we applied a digital low-pass filter (1^st^-order Butterworth filter; 6Hz cut-off frequency). Data were analyzed to provide estimates of reaction times, movement accuracies, maximal velocities and movement durations. Reaction time was operationalized as the temporal difference between the onset of the movement epoch and the moment when finger velocity exceeded a threshold of 11mm/s. Movement error sizes were characterized as the linear distance between the finger endpoint (calculated as average of the last five samples of the finger position during the REACH phase) and the border of the target circle.

### fMRI acquisition and analyses

MRI images were acquired using a 3T Siemens TRIO scanner using a twelve-channel head coil (Siemens, Ellwangen, Germany). For each subject, we obtained a T1-weighted magnetization-prepared rapid-acquisition gradient echo (MPRAGE) anatomical scan of the whole brain (176 slices, slice thickness: 1 mm, gap: 0 mm, in-plane voxel size: 1 x 1 mm, repetition time: 2300 ms, echo time: 2.92 ms, field of view: 256 x256, resolution: 256 x 256) as well as T2*-weighted gradient-echo planar imaging scans (EPI): slice thickness: 3.2 mm + 0.8 mm gap; in-plane voxel size: 3 x 3 mm; repetition time: 2000 ms; echo time: 30 ms; flip angle: 90°; field of view: 192 x 192 mm; resolution: 64 x 64 voxels; 32 axial slices. Overall, we obtained 2050 EPIs per subject in Experiment 1, which were collected during five consecutive runs. In Experiment 2 we collected again 2050 EPIs per subject over five runs. A single EPI volume completely covered the cerebral cortex as well as subcortical structures, apart from the most inferior aspects of the cerebellum which were not covered in several of our subjects. Functional data were processed using SPM8 (Wellcome Department of Cognitive Neurology, London, UK). In every subject, functional images were spatially aligned to the first volume in a series, and then coregistered to the T1 image. After that, a non-linear normalization of the structural image to a template in MNI space was performed. Parameters from normalization were then applied to the functional images. In the last step of data pre-processing, we smoothened all the functional images with a Gaussian filter of 6mm x 6mm x 8mm FWHM.

In subject-specific fMRI analyses we next specified a GLM for each individual including our four experimental conditions (“task” [DRT, CT] x “movement distance” [“NEAR”, “FAR”]). Each condition was modeled separately for each of our three trial epochs (CUE+MASK, DELAY, REACH). The regressor duration was defined according to respective epoch duration. The regressors were convolved with the canonical HRF-function of SPM8. Head motion parameters were included in the model as separate regressors. Fixation epochs weren’t explicitly modeled and served as an implicit baseline. To consider each subject’s individual functional brain organization, we detected planning areas significantly more active during the delay epoch in DRT than the respective epoch of CT trials in each subject (for that step, single subject activity maps were thresholded at p<0.001, uncorrected).

We additionally performed a group-level analysis to delineate the areas commonly activated by reach planning in our experiments. For this purpose we entered the respective (first level) contrast images in a second-level group analysis (one-tailed t test). In this step, we used a minimal cluster-size criterion (k>10 voxels) and a statistical threshold of p<0.001, uncorrected.

### Region of Interest Analyses

We used the results of the group-level analysis and anatomical landmarks (see below) to initially identify reach planning-related areas. Our ROI set consisted of two main areas: left dorsal premotor cortex located at the posterior end of the superior frontal sulcus, anteriorly to the hand area of M1 (PMd); the left posterior-medial portion of superior posterior lobule (SPL)^51^. The additional movement planning ROIs included were: the left anterior end of the intraparietal sulcus (aIPS); the left middle intraparietal sulcus (mIPS); and left supplementary motor area (SMA)^14^. For each of these ROIs and for each individual we next identified the coordinate of the voxel exhibiting the local maximum of the individual subject statistical contrast DRT>CT that was closest to the respective ROI group-coordinate. In addition we anatomically identified the hand area of left primary motor cortex^52^ due to its potential engagement in reach planning^7^ as well as left primary visual cortex (V1). The latter area served as a control for any activity related to visual stimulation, also because we are not aware of any findings showing its specific engagement in reach planning or execution. To avoid biasing our ROI-selection in individual subjects across both Experiments, as they were planning different movement types in each (circular [Exp. 1] vs. straight [Exp. 2]), in those subjects that participated in both of our experiments, we used the ROI coordinates of Experiment 1 also for Experiment 2 (5 out of 7 subjects) (compare fig. S2), For ROI analyses we always considered the average activity of voxels within a 3mm radius around the ROI center coordinate. Note that our ROI definition meets the criteria described by Kriegeskorte et al.^54^ to avoid circularity in data analysis.

### Time-resolved fMRI analysis

Using custom protocols written in Matlab (MathWorks) (compare ref. 33), we extracted and analyzed BOLD-signal timecourses for each of our ROIs. Importantly, we separately analyzed timecourses during the CUE and DELAY vs. the REACH epoch: timecourses for the CUE and DELAY phase were aligned to the onset of the CUE, and normalized to the baseline defined as a time window of -5s to -3s preceding CUE onset. As planning processes are likely to take place beginning as early as the presentation of the target cue we analyzed both trial epochs together. The signals for the REACH epoch were aligned to the onset of the REACH phase and normalized to the same baseline period as above. The timecourses were filtered with a digital high-pass filter (128s cutoff frequency) and interpolated at 1s temporal resolution (given the temporal jitter in our design).

To examine the effect of trajectory on the BOLD signal in each of the ROIs, we performed a time-resolved analysis of the timecourses with respect to their relative amplitude over the principle trial epochs (CUE/DELAY and REACH) using paired t-tests (compare “RESULTS” section). Only the significant differences spanning over three or more consecutive time points were taken into consideration. The comparison between the main ROIs was done with mixed-models ANOVA in R.

## AUTHOR CONTRIBUTIONS

AP: designed the paradigm, chiefly collected and analyzed the data, drafted and reworked the manuscript. AL: designed the paradigm, contributed to data collection and analysis and reworked the manuscript.

## ACKNOWLEDGMENTS

This work was supported by grants from DFG (CIN) and BMBF (FKZ 01GQ1002) to AL. We would like to thank Igor Kagan, Uwe Ilg, David J. Mack, Dan Arnstein, and all members of the NOD Lab for their valuable comments that helped us improve this work.

## FIGURE LEGENDS

**Figure S1.**
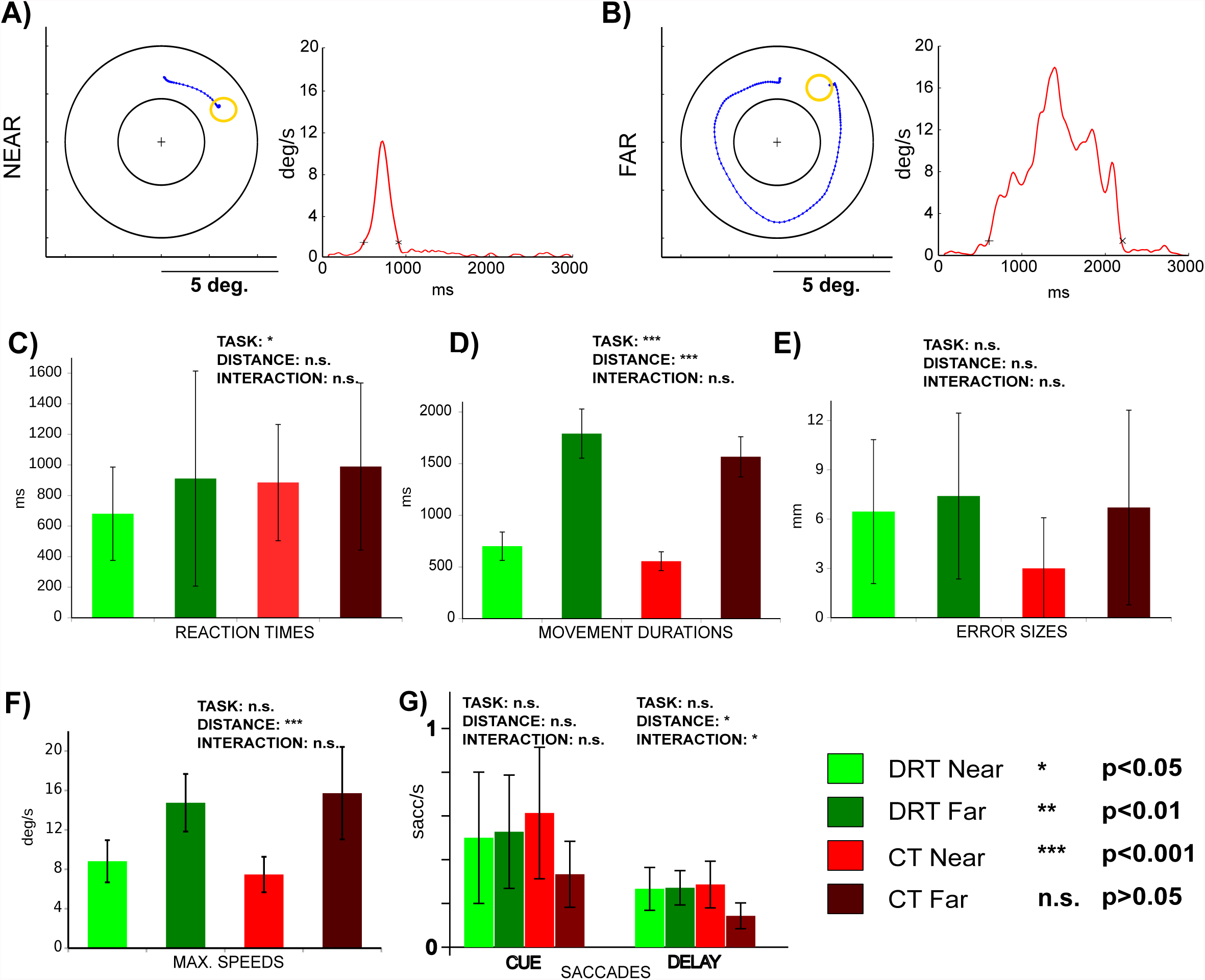
Movement performance in Experiment 1. A and B) Exemplary reach trajectories from a single subject (left panels) and the respective speed profiles throughout the REACH phase (right panels) for both a “NEAR” (F) and a “FAR” (G) condition are depicted. C-G) The individual panels show our estimates of behavioral performance as a function of “TASK” and “DISTANCE” and report the influence of these factors on the respective estimates as well as their interaction, as was assessed by two-way repeated measures ANOVAs (n.s. not significant; * p<0.05; ** p<0.01; *** p<0.001). C) Reaction times were significantly shorter in DRT than in CT. D) Movement durations were significantly longer in DRT than in CT and for “FAR” trajectories than “NEAR”. E) Error sizes were constant across all conditions. F) Maximal speeds were higher for “FAR” reaches. G) Average frequencies of fixational saccades in CUE an DELAY epochs of respective conditions. Saccades were less frequent in CT “FAR” than in all other conditions. Error bars represent SEM. See text for detailed statistics.

**Figure S2.**
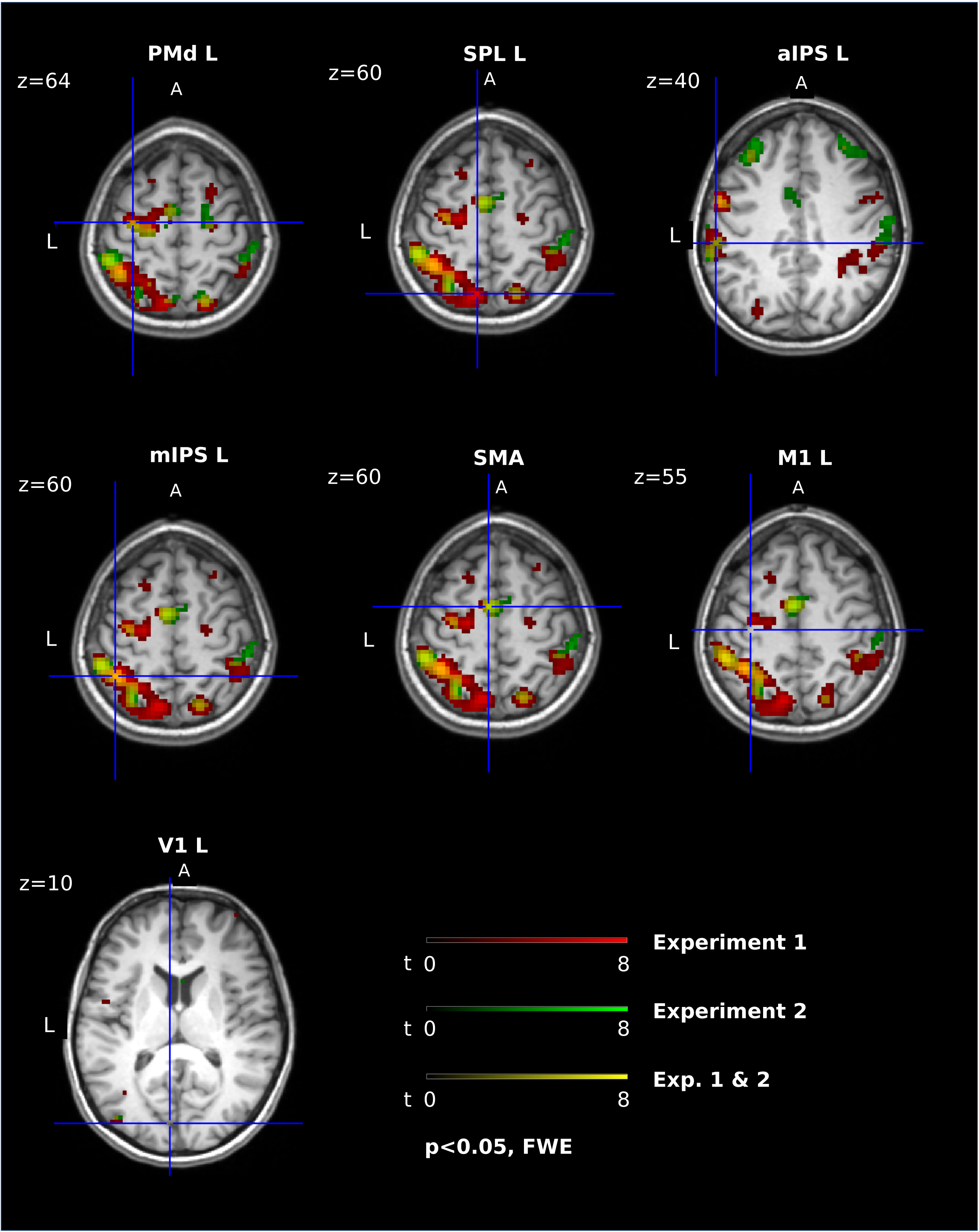
Comparison of planning regions recruited by our two experiments in a representative subject (DRT>CT; the overlaid maps of activity were thresholded at p<0.05, FWE-corrected for multiple comparisons). Red and green shaded regions denote clusters of planning activity specific to the delay phase of Experiments 1 and 2, respectively. Yellow shaded regions represents areas active in both experiments. Blue crosshairs indicate centers of clusters selected for subsequent ROI analyses (compare “METHODS”).

**Figure S3.**
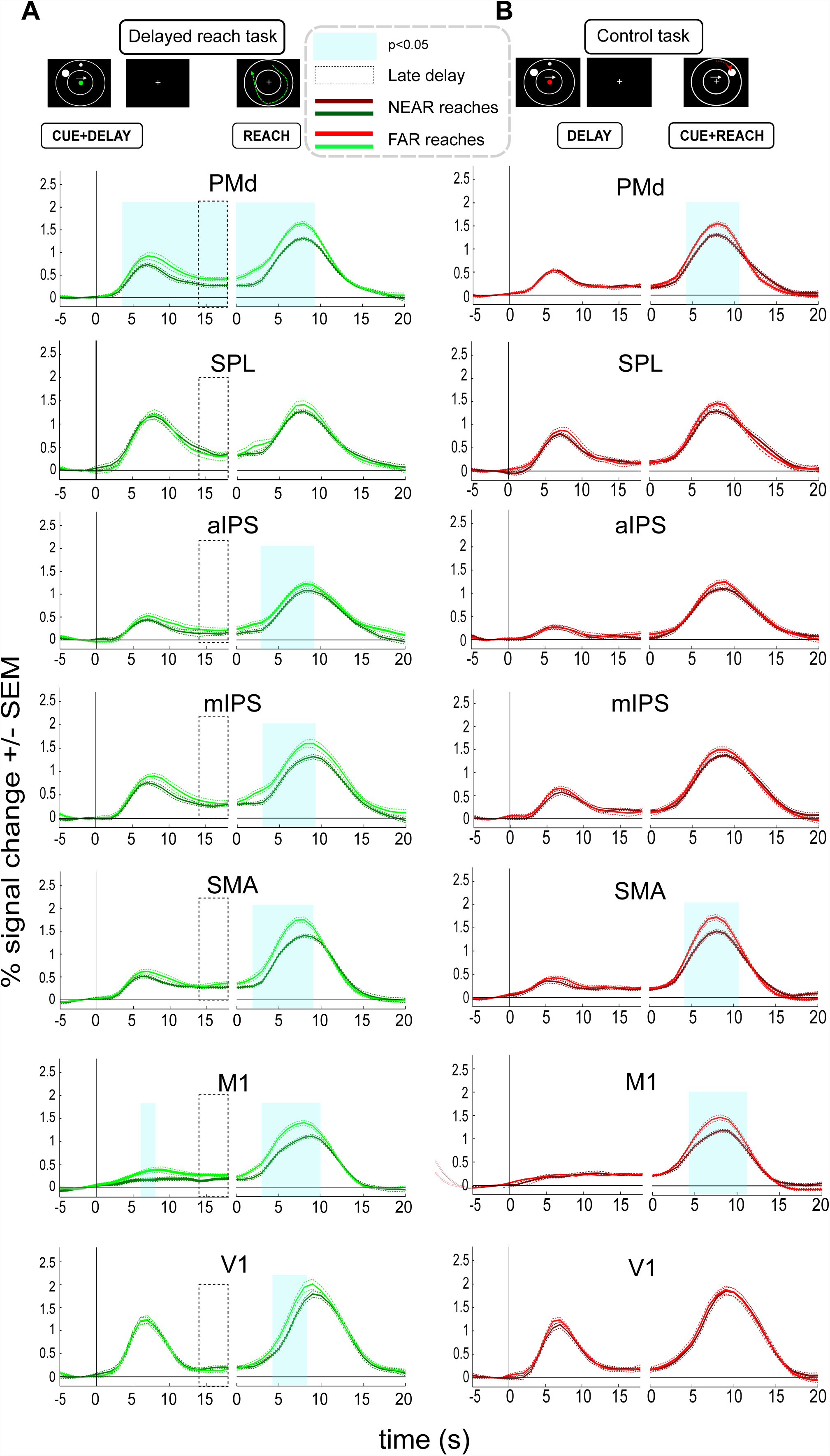
Timecourses of fMRI signals extracted from ROIs in the delayed reach (A) and control tasks (B) in Experiment 1. Left panels are aligned to CUE onset while right panels are aligned to REACH onset. Cyan-shaded areas represent time epochs during which paired t-test comparisons of signal amplitudes between “NEAR” and “FAR” reaches revealed statistically significant differences at p<0.05 for at least three neighboring time-points. Such differences were considered indicative of an influence of trajectory. In the DRT both PMd and M1 showed transient trajectory representation during the planning stage (leftward part of the panels, aligned to CUE phase onset). PMd, M1, SMA, mIPS, aIPS and V1 showed differences between the two types of trajectories during the reach stage (rightward part of the panels, aligned to REACH phase onset). **B)** No ROI shows planning-related differences in the CT. As in the DRT, PMd, SMA and M1 exhibit execution-related differences.

**Figure S4.**
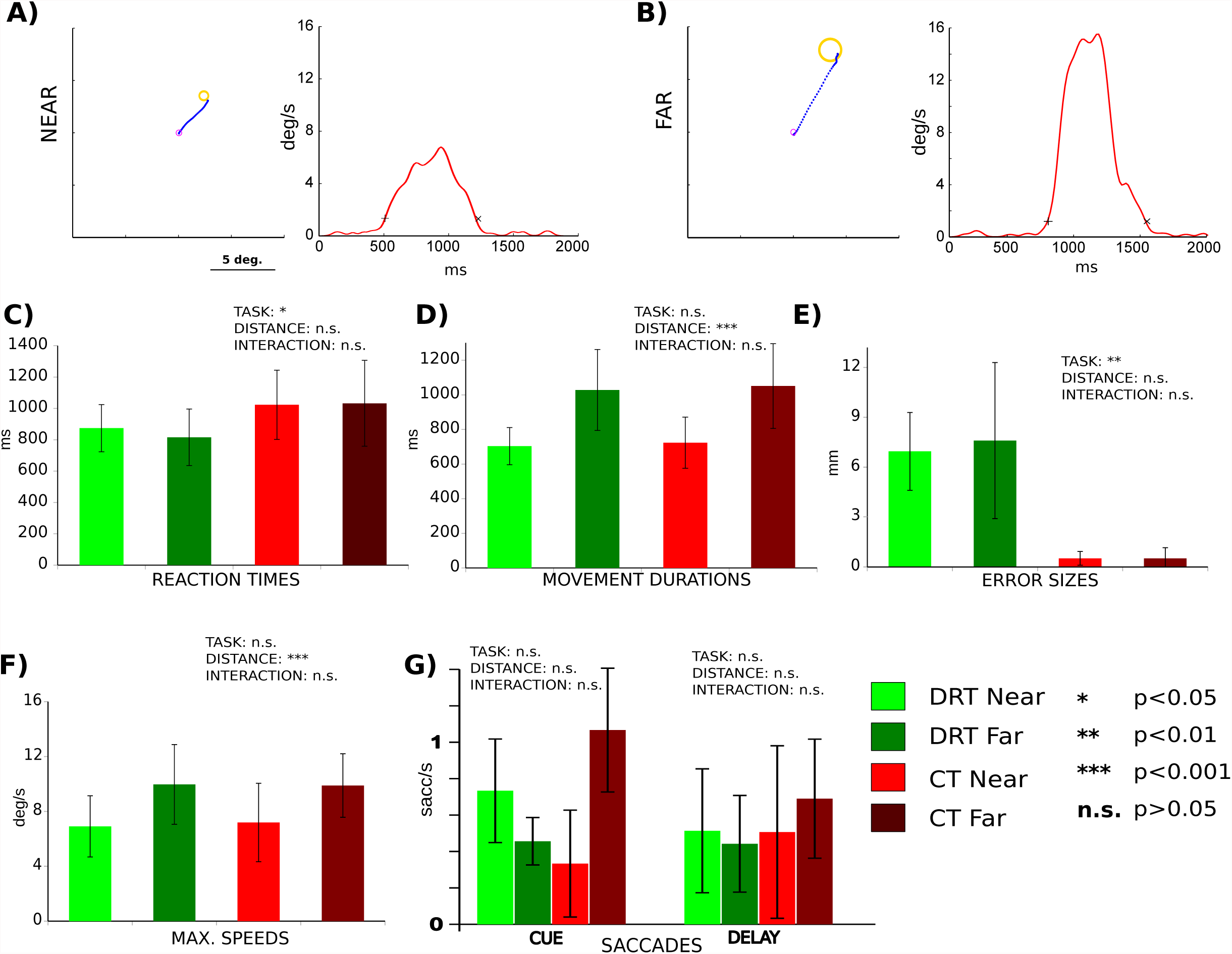
Movement performance in Experiment 2. A & B) Exemplary reach trajectories of a single subject (left panels) and the respective speed profiles (right panels) for both a “NEAR” (A) and a “FAR” (B) condition. C-G) The individual panels show our estimates of behavioral performance as a function of “TASK” and “DISTANCE” and report the influence of these factors on the respective estimates as well as their interaction, as was assessed by two-way repeated measures ANOVAs (n.s. not significant; * p<0.05; ** p<0.01; *** p<0.001). Error bars represent SEM. C) Reaction times were significantly shorter in DRT than in CT. D) Movement durations were significantly longer for “FAR” trajectories than “NEAR”. E) Error sizes were larger for DRT. F) Maximal speeds were higher for “FAR” reaches. See main text for detailed statistics. G) Average frequencies of fixational saccades in CUE an DELAY epochs of CT and DRT. The rates of fixational saccades were constant across all conditions.

**Figure S5.**
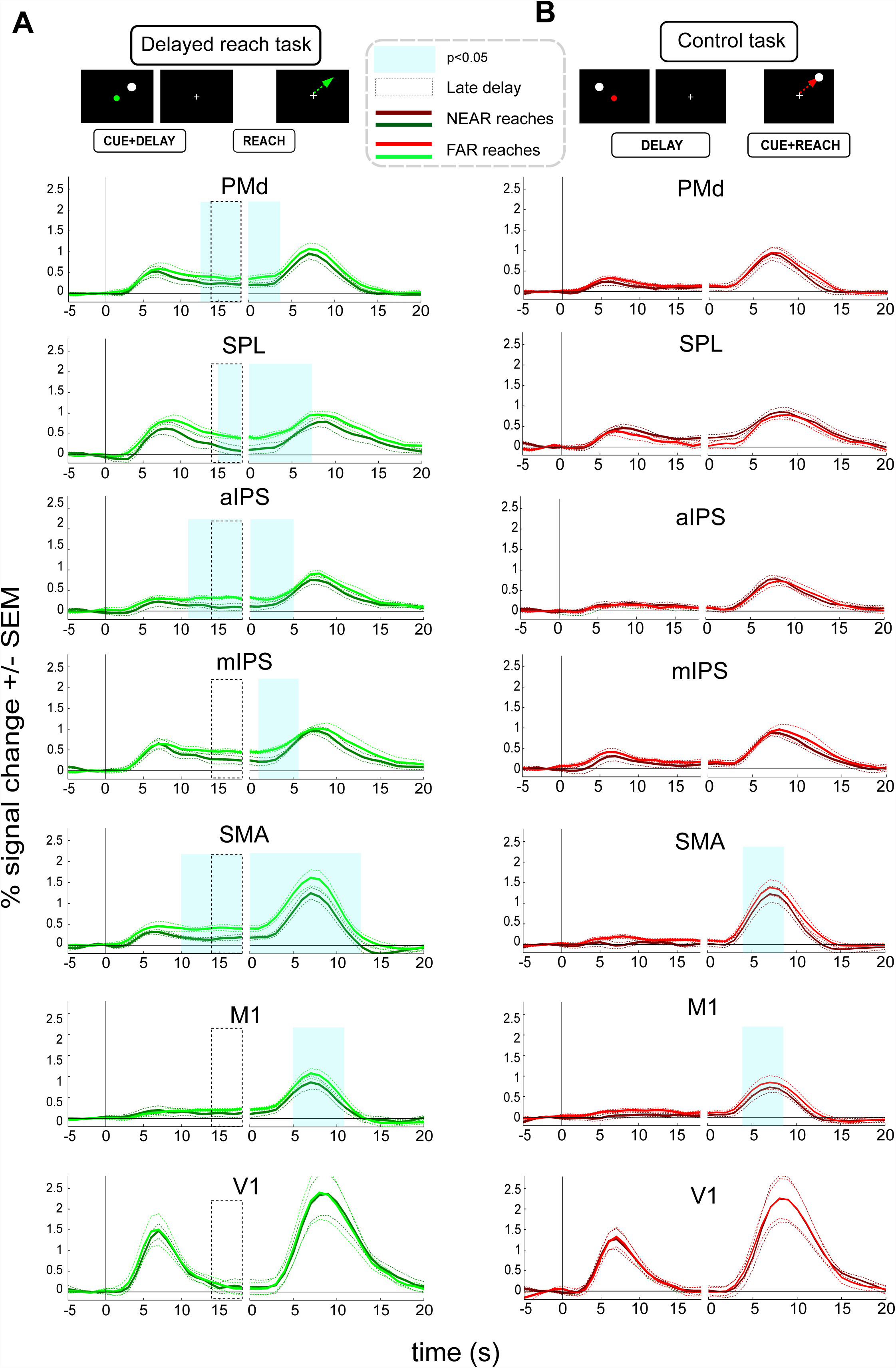
Timecourses of fMRI signals extracted from ROIs in the delayed reach and control tasks in Experiment 2. Left panels are aligned to CUE onset while right panels are aligned to REACH onset. Cyan-shaded areas represent time epochs during which paired t-test comparisons of signal amplitudes between “NEAR” and “FAR” reaches revealed statistically significant differences at p<0.05 for at least three neighboring time-points. A) PMd, SPL, SMA and aIPS show significant signal differences during the planning epoch in DRT. All areas except V1 show differences during the reach epoch. Note that some of these areas show differences only before the reach execution-related peak of the BOLD response, suggesting that some of these differences might still refer to planning during the late delay. B) In the control task, no ROI showed planning-related differences. However, both SMA and M1 exhibited differences during the reach stage (also compare to A and Figure S3).

